# Crystallographic fragment screening of the dengue virus polymerase reveals multiple binding sites for the development of non-nucleoside antiflavivirals

**DOI:** 10.1101/2025.03.31.646453

**Authors:** Manisha Saini, Jasmin C. Aschenbrenner, Francesc Xavier Ruiz, Ashima Chopra, Anu V. Chandran, Peter G. Marples, Blake H. Balcomb, Daren Fearon, Frank von Delft, Eddy Arnold

**Affiliations:** Center for Advanced Biotechnology and Medicine, Rutgers, the State University of New Jersey, Piscataway, NJ, 08854, USA; Department of Chemistry and Chemical Biology, Rutgers, the State University of New Jersey, Piscataway, NJ, 08854, USA; Diamond Light Source, Harwell Science and Innovation Campus, Fermi Ave, Didcot OX11 0DE, UK; Research Complex at Harwell, Harwell Science and Innovation Campus, Fermi Ave, Didcot, OX11 0FA, UK; Centre for Medicines Discovery, Nuffield Department of Medicine Research Building, Old Road Campus, Headington, Oxford, OX3 7FZ, UK

**Author notes:** These authors contributed equally to this work (Manisha Saini, Jasmin C. Aschenbrenner, Francesc Xavier Ruiz). Department of Biochemistry, Vanderbilt University School of Medicine, Nashville, TN, 37232, USA. Corresponding authors. (E.A.); (F.vD.); (D.F.); (F.X.R.).

## Abstract

Dengue viruses (DENV) infect approximately 400 million people each year, and there are currently no effective therapeutics available. To explore potential starting points for antiviral drug development, we conducted a large-scale crystallographic fragment screen targeting the RNA-dependent RNA polymerase (RdRp) domain of the non-structural protein 5 (NS5) from DENV serotype 2. Our screening, which involved 1,108 fragments, identified 60 hit compounds across various known binding sites, including the active site, N pocket, and RNA tunnel. Additionally, we discovered a novel binding site and a fragment-binding hotspot in thumb site II. These structural findings open amenable avenues for developing non-nucleoside inhibitors and offer valuable insights for future structure-based drug design aimed at DENV and other flaviviral RdRps.

## Introduction

Flaviviruses are single-stranded RNA viruses primarily transmitted by arthropods, causing severe illnesses in humans^1^. Over the past seven decades, the global spread and epidemic transmission of flaviviruses have been significant^2^. The mosquito-borne dengue viruses (DENV) infect approximately 400 million individuals annually, with over a quarter of the world’s population residing in endemic regions^1,3^. The disease symptoms range from self-limited dengue fever to severe dengue syndrome. The number of infections has steadily increased over the last 70 years, making DENV the most widespread arthropod-borne viral disease globally^3,4^. Strikingly, the World Health Organization has reported that around half of the world’s population is at risk of contracting dengue^5^. Any of the four DENV serotypes (DENV1 to 4) causes an acute febrile disease named dengue fever. However, many infected people, including a large proportion of children, develop life-threatening forms of the disease known as dengue hemorrhagic fever and dengue shock syndrome^6^. Prevention and control measures focus thus on vector management, as there is currently no specific treatment for dengue or severe dengue. Early detection and access to appropriate medical care are crucial in reducing fatality rates associated with severe dengue^1,7^.

The DENV viral particle consists of a positive RNA strand of 11 kb that forms the viral genome, including 5’ and 3’ untranslated regions (UTR) and a 5’ cap^8^. Upon infection, the RNA is translated into a single polypeptide chain within the endoplasmic reticulum (ER) membranes, which is then processed into ten proteins through proteolytic maturation by both viral and host cell proteases. The structural proteins, including the envelope, precursor-membrane, and capsid, encase the viral RNA. The non-structural (NS) proteins—NS1, NS2A, NS2B, NS3, NS4A, NS4B, and NS5— are expressed within the host cell but are not incorporated into the viral particle. With the assistance of host proteins, these NS proteins reorganize the internal structure of the cell, process the polyprotein to yield the mature viral proteins, replicate the viral RNA, and help the virus evade the immune system^9,10^.

NS5 is the largest and most conserved DENV protein, with approximately 100 kDa and with 70% sequence identity among the four dengue serotypes. It contains two key domains: a methyltransferase domain (MTase) at the N-terminus and RNA-dependent RNA polymerase (RdRp) at the C-terminus, connected by a 5–6 residue linker (residues 266–271) that plays a crucial role in determining NS5’s overall conformation and activity. The structure of NS5 is highly conserved across flaviviruses, suggesting the potential for designing broad-spectrum antiviral compounds targeting NS5^11^. However, there are different relative conformations between MTase and RdRp across the known full-length flaviviral NS5 structures^6^. The MTase domain (residues 1–265) is involved in capping the viral RNA and has guanylyl transferase, N7, and 2’-O ribose methylation activities. The RdRp domain is responsible for replicating the viral RNA, which is initiated *de novo* and facilitated by the “priming loop”. Upon transition to elongation, the priming loop retreats from the active site to allow elongation of the RNA duplex^12^. NS5 forms a larger replicase complex alongside the NS3 helicase, which assists in both polymerization and capping^13^. Beyond its role in viral genome replication, NS5 can also suppress the host immune interferon response, either through interaction with the signal transducer and activator of transcription 2 (STAT2) protein or by influencing RNA splicing within the host cell^9,14^. Recent structural work has provided improved detail on the conformational selection mechanism underlying STAT2 inhibition by NS5^15^.

Given that RdRp activity is absent in host cells, the RdRp domain of NS5 is a promising antiviral target for designing specific inhibitors with minimal on-target toxicity. Despite the crucial role of NS5 RdRp in the viral replication of DENV, the development of novel direct-acting antivirals has not exceeded the preclinical stage, except for the nucleoside analog AT-752 (Atea Pharmaceuticals), a chain terminator^6^ currently in clinical trials^16^. Despite this encouraging development and their historical success in drug development, it is known that nucleoside analogs have experienced a high attrition rate in clinical trials due to toxicity^17^. Regarding non-nucleoside inhibitors, through a structure-based fragment screening approach, researchers have developed allosteric inhibitors that target the so-called “N pocket” of DENV and Zika virus RdRps^18-21^. Nevertheless, retraction of the priming loop from the active site during enzyme elongation may alter the conformation of the N pocket, which may explain the weak affinities of the lead compounds developed despite the availability of multiple crystal structures of such protein-ligand complexes^19^.

The previous fragment screening campaigns (that yielded the N pocket inhibitor type) were performed using fragment cocktails against DENV3 RdRp. Here, leveraging the state-of-the-art XChem facility and pipelines^22^, we performed a large-scale crystallographic fragment screening campaign (1,108 fragments, with single fragment soaking) targeting DENV2 RdRp. Our study probed known binding sites (the active site, the N pocket, and the RNA tunnel site) and unveiled a fragment-binding hot spot (the thumb site II), providing insights for the development of novel DENV2 non-nucleoside inhibitors. Furthermore, these findings offer a framework for identifying and targeting analogous pockets across flaviviral RdRps.

## Results

### DENV2 RdRp crystallizes in a ligand-free form that diffracts to high resolution

The DENV2 NS5 RdRp apo crystal was solved with data diffracting to 1.56 Å resolution (PDB 7I2X). A single molecule was observed in the asymmetric unit (ASU) in the I222 space group, consistent with previously reported structures^19,23^. The structure closely aligns to the previous highest resolution apo (PDB 6IZY, 2.11 Å resolution) and ligand-bound (PDB 5K5M, 2.01 Å resolution) DENV2 RdRp structures (RMSD = 0.385 Å and RMSD = 0.787 Å, respectively). Despite the improved resolution, the apo structure presented similarly disordered regions as the previous (Fig. S1, Supplementary Data), including a disordered priming loop (similarly to PDB 6IZY and conversely to PDB 5K5M, with a small molecule bound in the N pocket) and an ordered motif G (similarly to PDB 6IZY and conversely to PDB 5K5M). Preliminary screening of this crystal form with a small subset of fragments validated the suitability for large-scale crystallographic fragment screening.

### Crystallographic fragment screening comprehensively maps many binding sites in DENV2 RdRp

We soaked 1,108 fragments into the aforementioned RdRp apo crystals. Soaking of crystals with 10% DMSO for 1-3 h showed minor effect on diffraction quality, yielding 918 datasets with resolution between 1.56 and 2.43 Å (Fig. 1A). Notably, ∼60% of these datasets diffracted to a resolution better than 2.0 Å, with an average resolution of 2.1 Å overall. After thorough refinement of the ground-state model, the Pan-Dataset Density Analysis (PanDDA)^24,25^ and PanDDA 2^24^ pipelines were run, identifying a total of 35 hits. Additionally, given the presence of several disordered loops and relative crystal heterogeneity, we ran the Cluster4x algorithm^26^. This tool analyzes and clusters multi-dataset crystallographic data, grouping similar datasets together based on reciprocal lattice intensities or C-alpha atom positions, thereby improving the signal-to-noise ratio. PanDDA2 was then run on these pre-clustered datasets which enabled the identification of an additional 25 hits, a 73% improvement. Following meticulous modeling, data curation, and cross-reviewing, a total of 60 fragment-bound structures (with 69 binding events, with 7 fragments bound in two sites and 1 fragment in three sites) and one ground-state structure of DENV2 NS5 RdRp were validated and subsequently deposited in the Protein Data Bank (PDB). PDB identifiers, names, ligand descriptors, and data collection and refinement statistics are available in Table S1.

**Figure 1.**
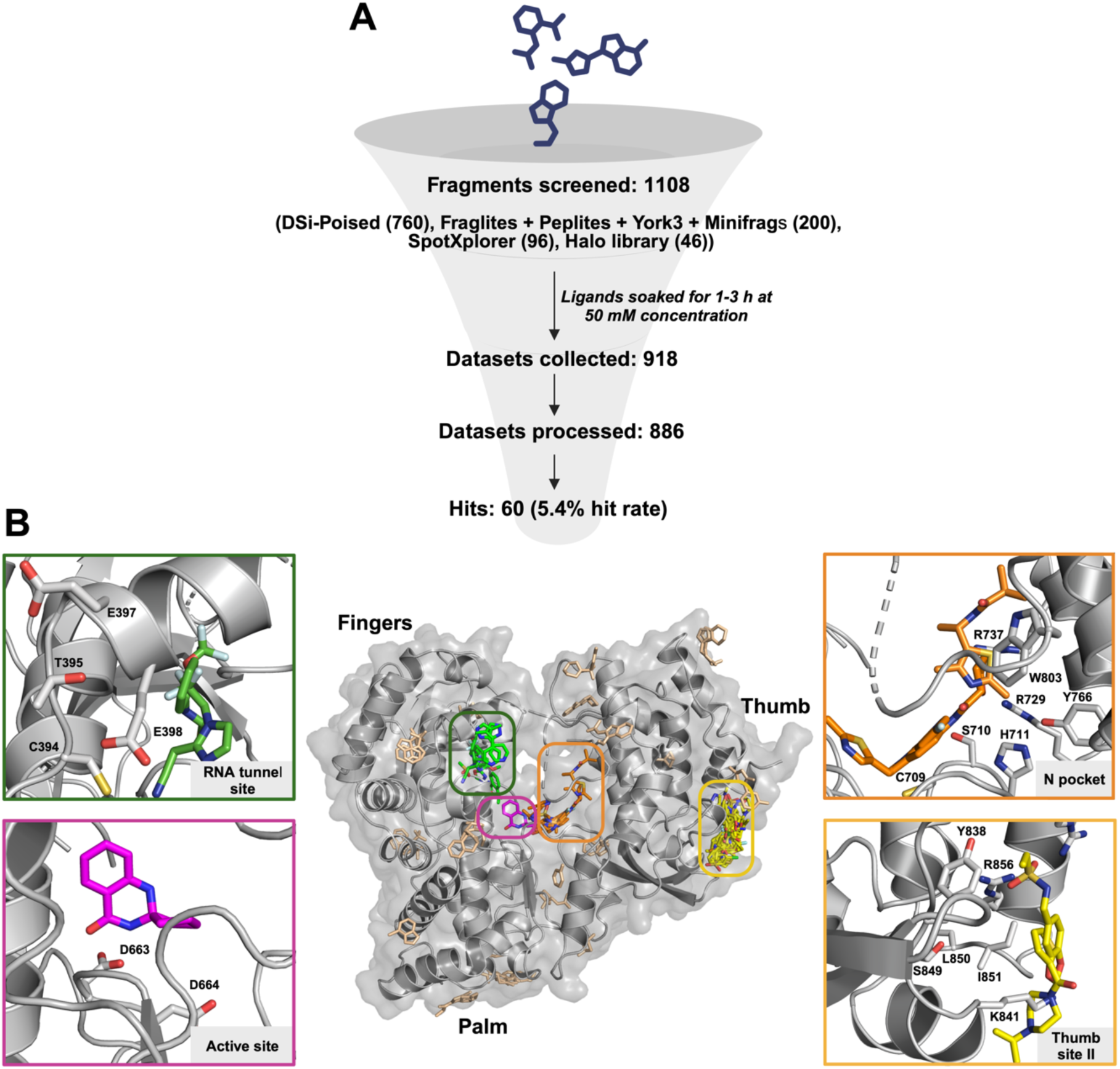
Bound fragments comprehensively sample the whole DENV2 RdRp. A) Overview of the fragment screening campaign presented in this study. B) The crystallographic fragment screen reveals multiple binding sites: we selected four sites that are highlighted in green, pink, orange and yellow boxes. The green box highlights binders in the RNA tunnel site, the orange box showcases N pocket (or primer grip site) binders, the pink box indicates the single active site binder found, and the yellow box shows binders for the novel thumb site II. The remaining fragments are displayed in light brown. All the fragment hits are depicted as sticks. The protein backbone is represented as grey cartoon.

We identified multiple hits away from the active site, many on the surface of the protein (Fig. 1B). However, it remains a challenge to rationalize and validate the putative allosteric binding pockets in the absence of fragment-originated small molecules that can be assayed in biochemical inhibition or binding assays^27^. Based on published structures of RdRps from the *Flaviviridae* family deposited in the PDB (e.g., RdRps from flaviviruses as DENV, Zika virus; and from hepaciviruses as hepatitis C virus, HCV), we have nominated in the current study four sites with the potential of driving development of non-nucleoside small molecule inhibitors: one active site^13,15,28^, and three allosteric sites; N pocket (or primer grip site)^19,23,29^, RNA tunnel site^30-32^, and thumb site II ^33,34^ (Fig. 1B). Fig. S2 displays all hits grouped in the previous four top sites.

### A fragment hit binds in the DENV2 RdRp active site, recapitulating the initiating ATP

RdRps belong to the template-dependent nucleic acid polymerase superfamily. While their sequence and length are variable, they contain seven conserved motifs (from A to G) essential for nucleotide selection and catalysis. Each of the motifs adopt a specific and conserved fold and are encompassed into one of three subdomains: fingers, palm and thumb, in analogy to the polymerase domain’s resemblance to a cupped right hand^10^. The palm domain spans residues 497–542 and 601–705 (Fig. S1), comprising a small antiparallel β-strand platform (β7 and β8) surrounded by eight helices^1,10,29^. The domain is the most structurally conserved region among all known polymerases. The catalytic active site in motif C—with the strictly conserved residues GDD (G662, D663 and D664 in DENV2)—is located in the turn between strands β7 and β8 (Fig. S1^23,35^).

We identified one fragment (Z48978335) in the DENV2 RdRp active site cleft (Fig. 2A). The D663 backbone NH and the N610 side chain are hydrogen-bonded to the carbonyl of the fragment, while Y607 is engaged in a T-shaped π-π stacking interaction with the pyrimidine ring of Z48978335. The amide NH in the dihydroquinazolinone moiety of the fragment also displays a hydrogen bond with the oxygen atom of the carboxylate of D663 (Fig. 2B-C). A similar positioning is observed for ATP bound to the related Japanese Encephalitis Virus (JEV) RdRp (PDB 4HDH), which might be the initiating nucleotide for the *de novo* synthesis in flaviviral RdRps^36^. ATP forms a hydrogen bond with D668, residue that is analogous to D664 in DENV2 RdRp^37^ (Fig. 2D). A superimposition of the fragment (magenta) and ATP (cyan) within the same binding pocket (Fig. 2E) highlights the fragment’s binding position relative to the naturally occurring substrate.

**Figure 2.**
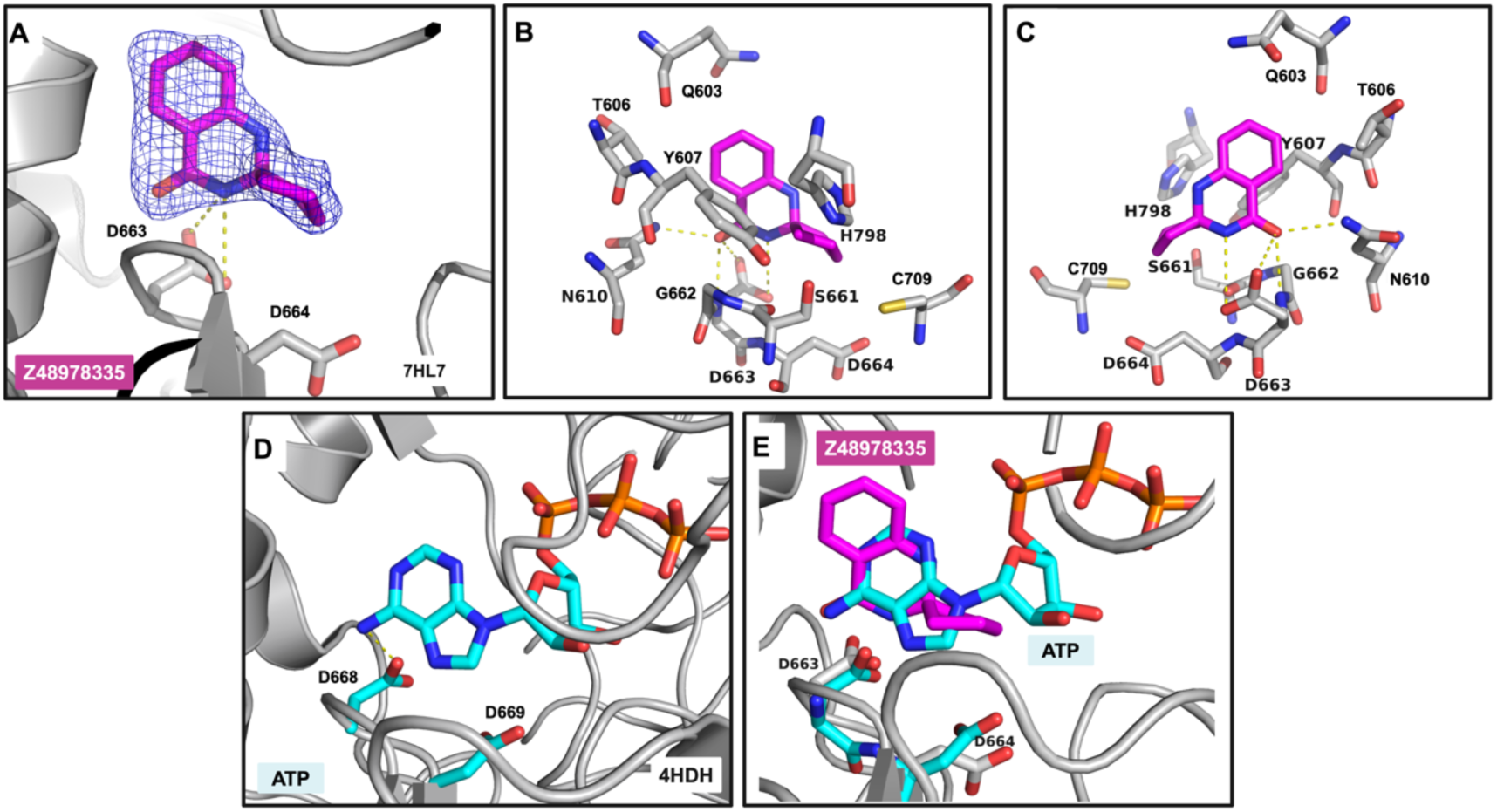
Fragment binding in the active site cleft. A) Fragment binding in the active site cleft of DENV2 RdRp (PDB 7HL7). The bound fragment is shown with pink sticks surrounded by a blue mesh corresponding to the PanDDA event map at 1σ contour. B) shows the residues surrounding Z48978335, C) is analogous to B) with a 180° rotation in the x-axis. Polar interactions in panels A to C are displayed as yellow dashed lines. D) illustrates the binding of ATP bound to the Japanese encephalitis virus (JEV) RdRp (PDB 4HDH) forming a hydrogen bond (yellow dashed line) with D668, which is analogous to D663 of DENV2 RdRp. E) Superimposition of the bound ATP to the JEV RdRp onto the DENV2 RdRp complexed to Z48978335.

### The previously described N pocket is a hot spot for fragment binding

This site was designated as the “N pocket” by Novartis scientists in a previous fragment screening campaign^20^ and is in a similar location to the Palm I in HCV NS5B^33,34^. It is a binding cleft between the palm and the thumb subdomains, with the “exit dsRNA loop” (residues 506-513, Fig. S1) in the front, the primer grip (residues 707-712, Fig. S1) in the back, and the priming loop (residues 789-805, Fig. S1) closing the pocket on the top (front view, Fig. 3A and 3C). This pocket is important for NS5 polymerase *de novo* initiation activity and viral replication^6,19^. It is also known as motif E or “primer grip” (as it comprises the later), is conserved across RNA viruses and retroviruses, and may be involved in correct positioning of the priming nucleotide ATP during initiation and the primer terminus during elongation^38-40^. The literature shows that N pocket binders provide good initiation inhibition but slightly poorer inhibition of elongation^18-21^.

**Figure 3.**
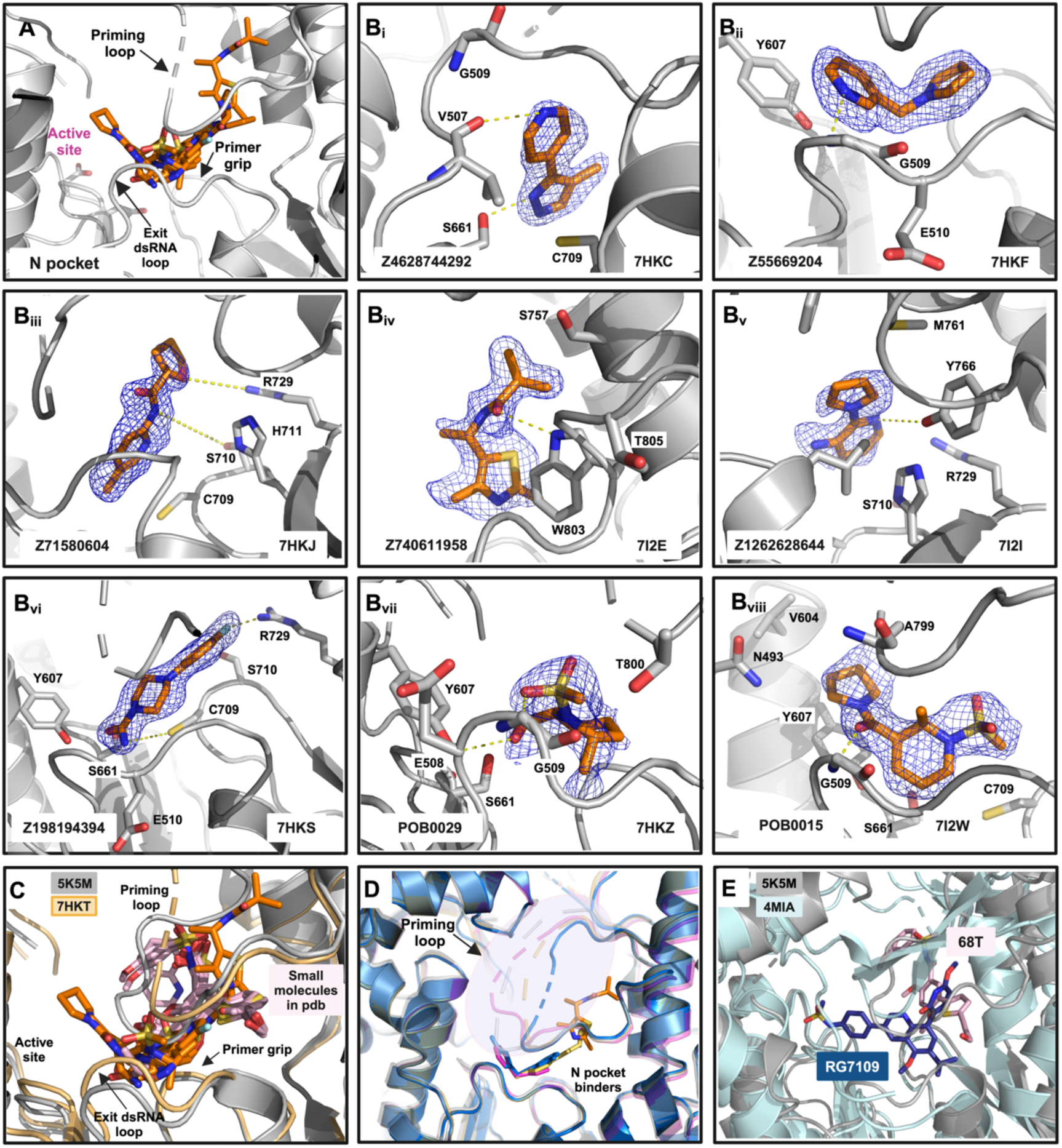
Fragments binding to the N pocket. A) Superimposition of the N pocket binders and protein elements surrounding them. B(_i-vii_) Each sub-panel (Bi–Bviii) represents distinct fragments (e.g., Z4628742292, Z55669204) bound in different orientations within the N pocket. The fragments are shown with orange sticks and the PanDDA event map (at 1σ contour) is shown as a blue mesh. C) Superimposition of previously reported small molecules from DENV 3/2 structures (PDB 5F3T, 5F3Z, 5F41, 5HMW, 5HMX, 5HMY, 5HMZ, 5HN0, 5I3P, 5I3Q, 5K5M, 6H9R, 6H80 and 6IZX) and current fragment screening hits in the same orientation (fragments in orange, small molecules in light pink). Polar interactions are displayed as yellow dashed lines. D) Ensemble of conformations that the priming loop is adopting in fragment-bound datasets (PDB 7HKF (pink), 7HKJ (yellow), 7I2E (grey cartoon with orange sticks) and 7HKT (blue). E) Superposition of DENV2 RdRp N pocket inhibitor 68T (PDB 5K5M) with HCV NS5B RdRp with RG7109 (PDB 4MIA), which is a close analog of the FDA-approved drug dasabuvir.

We identified a total of nine fragment hits that bind to the N pocket (Fig. 3B_i__-viii_). The complete list of fragments is provided in Fig. S2. As previously commented, the priming loop is disordered in DENV RdRp apo structures, but it is ordered in the presence of N pocket small molecule binders, as they occupy the N pocket in the interface between the primer grip and the priming loop (Fig. 3C). Our fragment hits, on the other hand, mainly form interactions with residues in the area in between the exit dsRNA loop and the primer grip, and do not display significant interactions with the priming loop. As a result, the priming loop in our liganded structures is partially disordered and is showing an ensemble of conformations (Fig. 3D). Fig. 3C portrays the superimposition of the current fragment hits and the previously reported small molecules in complex with DENV2/3 RdRp^19-21,41^. We discuss in the “Fragment progression” section the prospects for leveraging all this chemical matter.

### Fragment hits in the conserved RNA tunnel site could interfere with RNA template binding

The RNA tunnel site is a hydrophobic pocket surrounded by helices α5 (preceding motif G, Fig. S1) and α8-α9 (immediately after motif F, Fig. S1 and Fig. 4A), with two flexible loops (motif G or pinky, and the loop before motif B, Fig. S1) that move depending on fragment binding (Fig. S3A). This binding site, involving the conserved motifs B, G and F, was previously observed in structures of the DENV3 RdRp with inhibitors NITD107 (PDB 3VWS)^42^, HeE1-2Tyr (PDB 5IQ6)^43^, and NITD-434 (PDB 6XD0)^32^. A total of five binders were identified at this site (Fig. 4C, D(_i-v_)). The fragment hits mainly display hydrophobic interactions with F399 and F486, and polar interactions, i.e., H-bond with sidechains of E494, and with the backbone of Y607 and V604 (Fig. 7B(_i-v_) and S3B). As this site comprises the pinky finger region or motif G that interacts with the template RNA and plays an important role in restricting the template strand movement, the fragment hits could inhibit or alter the binding and positioning of the RNA template, which is essential for the start of RNA replication, either by direct competition or by altering the pinky conformation (Fig. 4E). The literature demonstrates that RNA tunnel binders provide good initiation inhibition and slightly poorer inhibition of elongation^32^.

**Figure 4.**
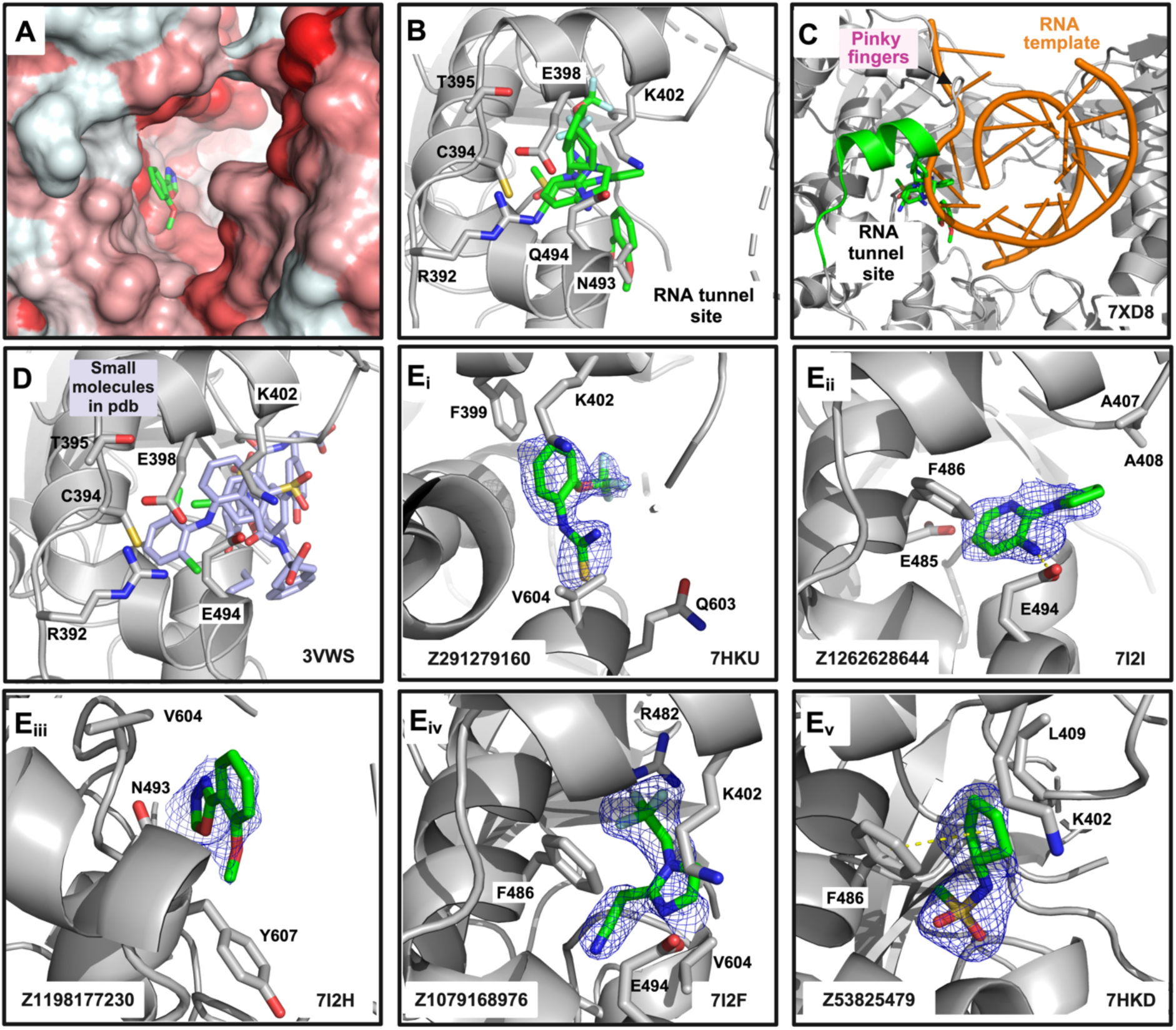
Fragment binding to RNA tunnel site. A) Surface representation of the RNA tunnel site, colored according to the hydrophobicity scale^44^ (red correspond to the most hydrophobic atoms, while white to the less hydrophobic atoms). B) Superimposition of the RNA tunnel site binders and protein elements surrounding them. C) Positioning of the RNA tunnel site and its binders near the RNA template (orange) and the pinky fingers. The protein/RNA structure shown corresponds to PDB 7XD8. D) Small molecules from previously solved structures (PDB 3VWS and 5IQ6) occupy the same site, underscoring its ligandability. The protein structure shown corresponds to PDB 3VWS. E) Each sub-panel (i-v) represents distinct fragments bound in different orientations within the RNA tunnel site. The fragments are shown in green sticks with the PanDDA event map at 1σ contour as a blue mesh.

### The “thumb site II”—a validated site in HCV RdRp—is DENV2 RdRp’s “hottest” spot

A total of twenty-nine binders were identified at “thumb site II” (Fig. 5A, 1B). Eight representative hits are shown in Fig. 5B(_i-viii_). The complete list of fragments is provided in Fig. S2. Each panel highlights different fragments (yellow sticks) bound to the thumb site II. Nevertheless, a word of caution is needed, as many hits feature at least one interaction that may be mediated by crystal packing, e.g., R698 of the neighboring asymmetric unit (Fig. S4A). Along these lines, Fig. 5B displays hits that seem to make “genuine interactions” (i.e. independent from crystal packing) with thumb site II. The predominant interactions observed are polar interactions with R770, R856, K887 and R888. Indeed, the thumb site II contains an arginine patch involved in RNA recognition, which, if the interaction is disrupted, results in loss of viral replication, as it occurs for mutation to alanine of the mentioned R770 and R856^45^.

**Figure 5.**
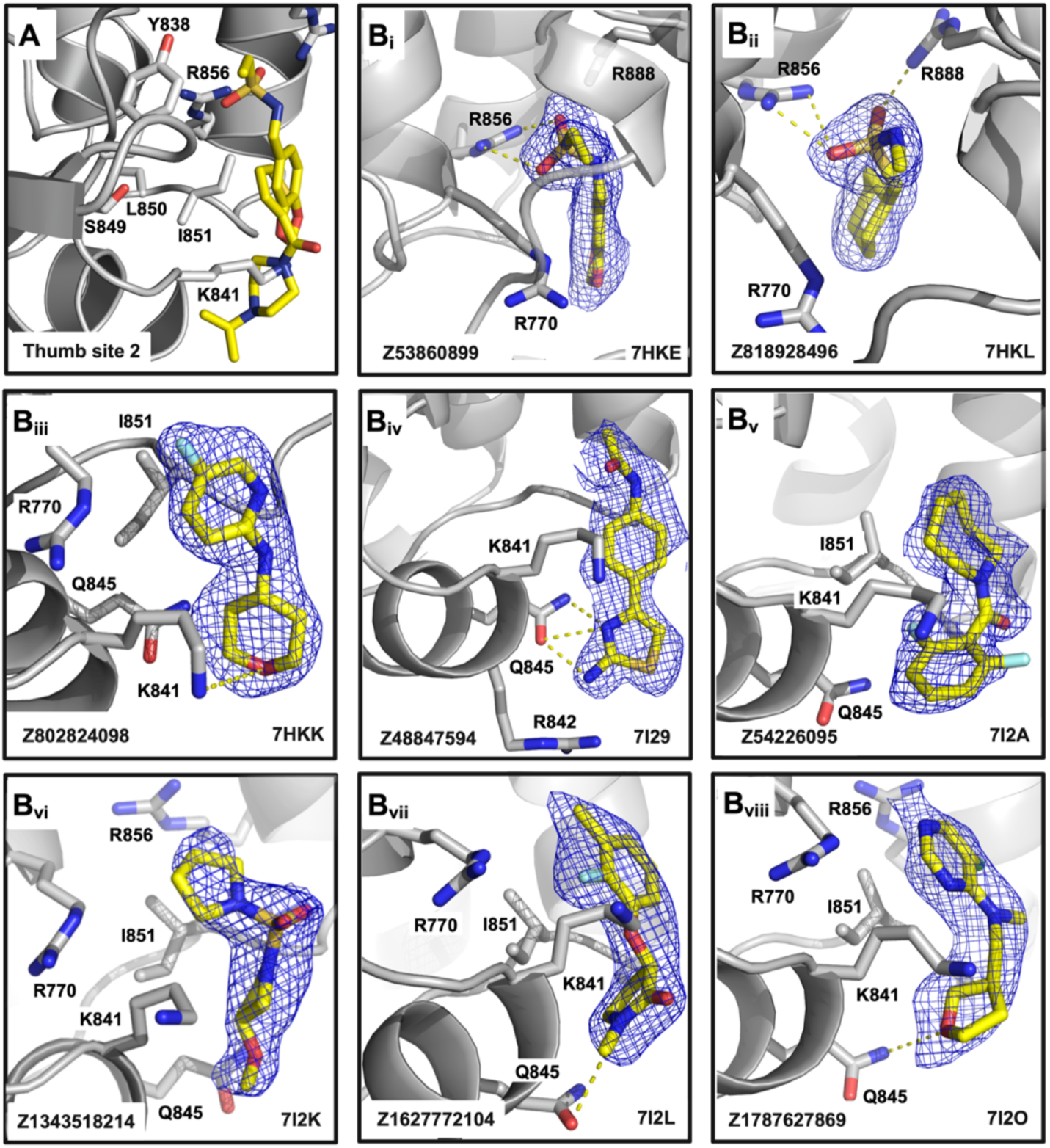
Fragments binding to the thumb site II. A) Overview of the thumb site II; showing key residues of the protein backbone interacting with bound fragments. B(_i-vii_) Each panel represents a detailed view of individual fragments (yellow sticks) bound to the thumb site II, with the corresponding PanDDA event maps at 1σ contour depicted as blue meshes. Key interactions include hydrogen bonds (yellow dashed lines) with residues such as R770, Q845, K841, and R856, as well as hydrophobic interactions with residues like I851 and L850.

The “thumb site II”, which we named based on the similar location to the analogous pocket in HCV RdRp (Fig. 6A) is located on the thumb subdomain near the C-terminus of NS5. This site has been heavily targeted in HCV drug discovery campaigns, with inhibitors such as filibuvir^47^, lomibuvir^47^, and radalbuvir^47^ (Fig. S4B) having reached phase II clinical trials. The thumb site II seems to prevent structural rearrangements in the HCV RdRp-RNA complex required for the transition from initiation to elongation^33,34^. Additionally, while their binding to the thumb site II was not acknowledged^21^, two N pocket binders solved in complex with DENV3 RdRp (PDB 5HMW, 5HMX) also had a second binding site coinciding with the “thumb site II” of DENV2 RdRp (Fig. 6B).

**Figure 6.**
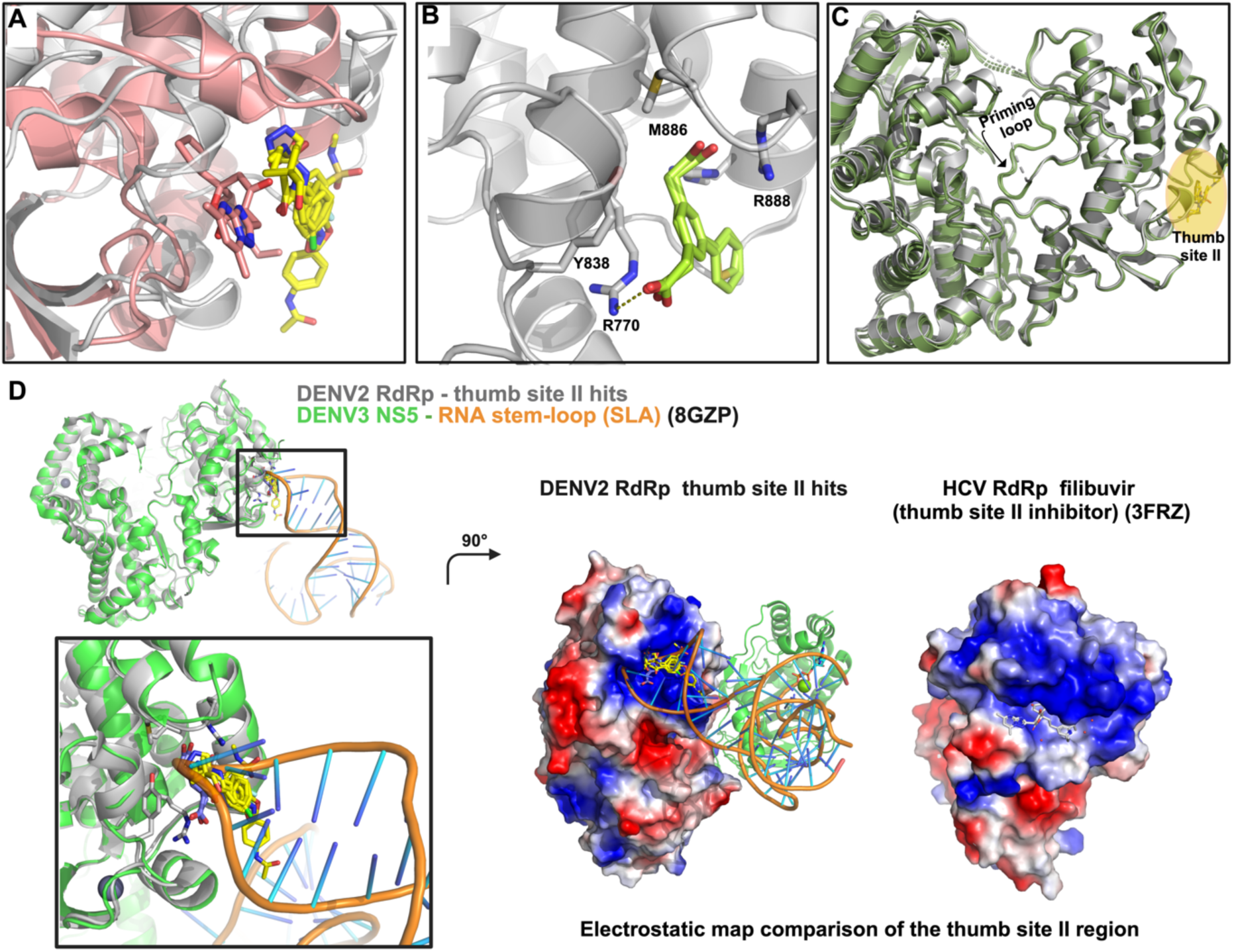
Mechanistic insights into thumb site II hit binding and function. A) Superposition of the thumb site II hits (yellow sticks) in DENV2 RdRp (grey cartoon) with the HCV NS5B RdRp thumb site inhibitor II inhibitor filibuvir (or PF868554, PDB 3FRZ). Both protein and compound are displayed in light pink. B) Thumb site II (light green sticks) binding site of the co-crystallized hits with DENV3 RdRp (in grey cartoon and sticks, PDB 5HMW, 5HMX). C) Comparison of the ground state (or apo) structure of DENV2 RdRp (grey cartoon, PDB 7I2X) with the DENV2 RdRp (in green cartoon) structures with fragments Z1343518214, Z32327641 and Z32665176 (in yellow sticks, PDB 7I2K, 7I2T and 7I2U), highlighting the ordered priming loop of the later. D) Superposition of the DENV2 RdRp-thumb site II hits and the DENV3 NS5 complex with the RNA stem loop (SLA, PDB 8GZP) as indicated in the legend, focusing on the thumb site II binding site. To note that a recent preprint^46^ and deposited structure of the DENV2 NS5/SLA complex (PDB 9DTT) appeared during the preparation of this manuscript. Additionally, the electrostatic potential surfaces of the indicated proteins are displayed to showcase the positively charged nature of both thumb site II binding sites.

### Fragment Progression

The fragment hits found in this campaign show several opportunities for progression to small molecule inhibitors of DENV2 RdRp. We have summarized these in Fig. 7.

**Figure 7.**
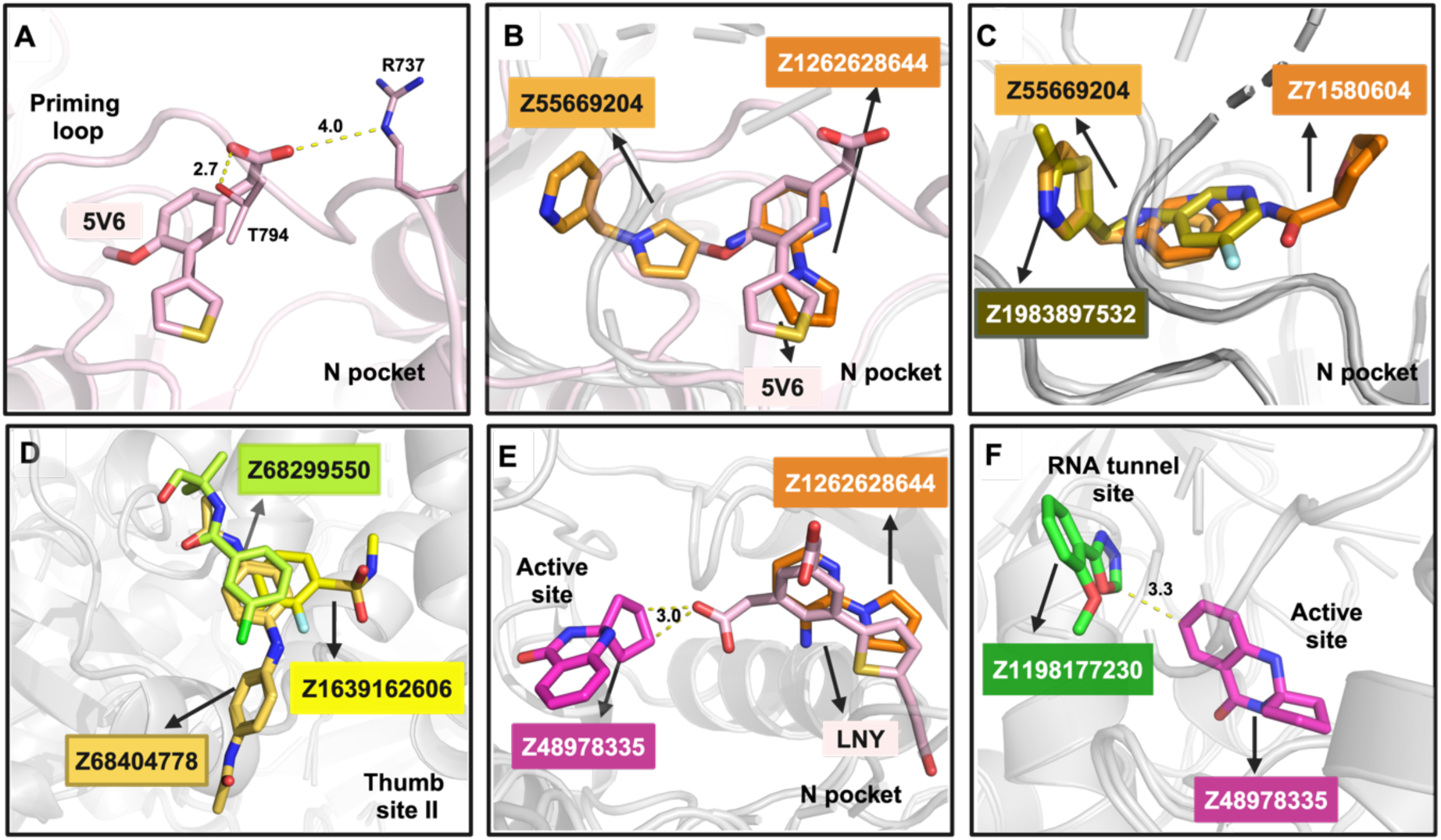
Fragment progression prospects derived from the fragment hits. A) The acetic acid moiety of DENV3 RdRp fragment hit 5V6 (light pink, PDB 5F41) presumably stabilizes the priming loop through polar contacts with residues T794 (main chain) and R737 (side chain). B) and C) N pocket fragment merge prospects originating from superposition of the indicated DENV2 RdRp-bound structures. D) Thumb site II fragment merge prospects originating from superposition of the indicated DENV2 RdRp-bound structures. Fragment linking prospects between the active site hit Z48978335 and E) indicated N pocket hits or F) indicated RNA tunnel hit, originating from superposition of the indicated DENV RdRp-bound structures (LNY is a fragment bound to a DENV3 RdRp complex, PDB 5HMY). Minimal distance between fragments to be linked in E) and F) are indicated.

Regarding the N pocket, none of our fragment hits stabilize the priming loop (Fig. 3A and D). Comparison with the fragment hits found in DENV3 RdRp^20^ shows that Z55669204 has a common core to the biheterocyclic core of the former. However, the DENV3 fragments have an additional acetic acid moiety that presumably stabilizes the priming loop (Fig. 7A). To note that further elaboration of this fragment scaffold replaced the acetic acid moiety with an acyl-sulfonamide bioisostere^19^. Fig. 7B shows a fragment merge of the biheterocyclic core of Z1262628644 (with the additional acetic acid moiety) and with Z55669204. Such a small molecule would keep the priming loop ordered while extending further towards the exit dsRNA loop. Alternatively, it may be worthwhile to attempt fragment merges with only the current fragment hits, providing a totally novel chemical scaffold (Fig. 7C). Regarding the thumb site II, as mentioned, we prioritize the hits with “genuine” interactions (i.e., main interactions are not due to crystal packing) for progression, illustrated with the merge of Z68299550, Z68404778 and Z1639162606 (Fig. 7D).

Another appealing avenue is linking fragment hits of neighboring sites. Concretely, we envision linking the active site hit, Z48978335, with hits of the: 1) the N pocket, and 2) the RNA tunnel site. Regarding 1), we hypothesize that leveraging the second acetic acid moiety of “compound 15” (developed from the aforementioned biheterocyclic core fragment found in the previous DENV3 RdRp Novartis screen^21^) allow to trace a feasible linking path (Fig. 7E). Meanwhile, the active site hit could also be linked to the most proximal hit of the RNA tunnel site, Z1198177230 (Fig. 7F).

Finally, the abundance of the hits obtained in these sites will allow to utilize algorithmic design of fragment merges for the three sites, as we and collaborators have done before^48^. This approach may be especially useful in the case of the RNA tunnel site, with no straightforward manual merges, and to leverage the 29 hits of the thumb site II, especially for the hits with a relevant crystal packing interaction component.

## Discussion

Despite the exciting progress initiated by Janssen’s development of JNJ-1802, a first-in-class antiviral agent against dengue^2^, there is still a lack of therapeutic treatments that effectively combat dengue as well as most other flaviviral diseases^6^. Therefore, our current work focused on DENV2 RdRp, one of the most conserved targets with potential for broad-spectrum antiviral activity against flaviviruses, is well aligned with the need for pandemic preparedness collectively^49,50^.

Thus, the 60 hits identified by this crystallographic fragment screen provide novel chemical matter for the development of non-nucleoside small-molecule antivirals that target the DENV2 as well as other flaviviral RdRps. Our study has identified four distinct sites, which are presumed to have ligandability, as they have been characterized in related systems (especially HCV RdRp) over the past years (Fig. 8). The structural data from our fragment screen are accessible through the Fragalysis web tool (see Data availability section), aligned in a common reference to an analogous Zika virus NS5 RdRp fragment screen and other relevant flaviviral RdRp-ligand structures, allowing their seamless and interactive exploration. These combined efforts may help pave the way for the development of non-nucleoside pan-flaviviral antivirals. We will discuss next some prospects to be pursued.

**Figure 8.**
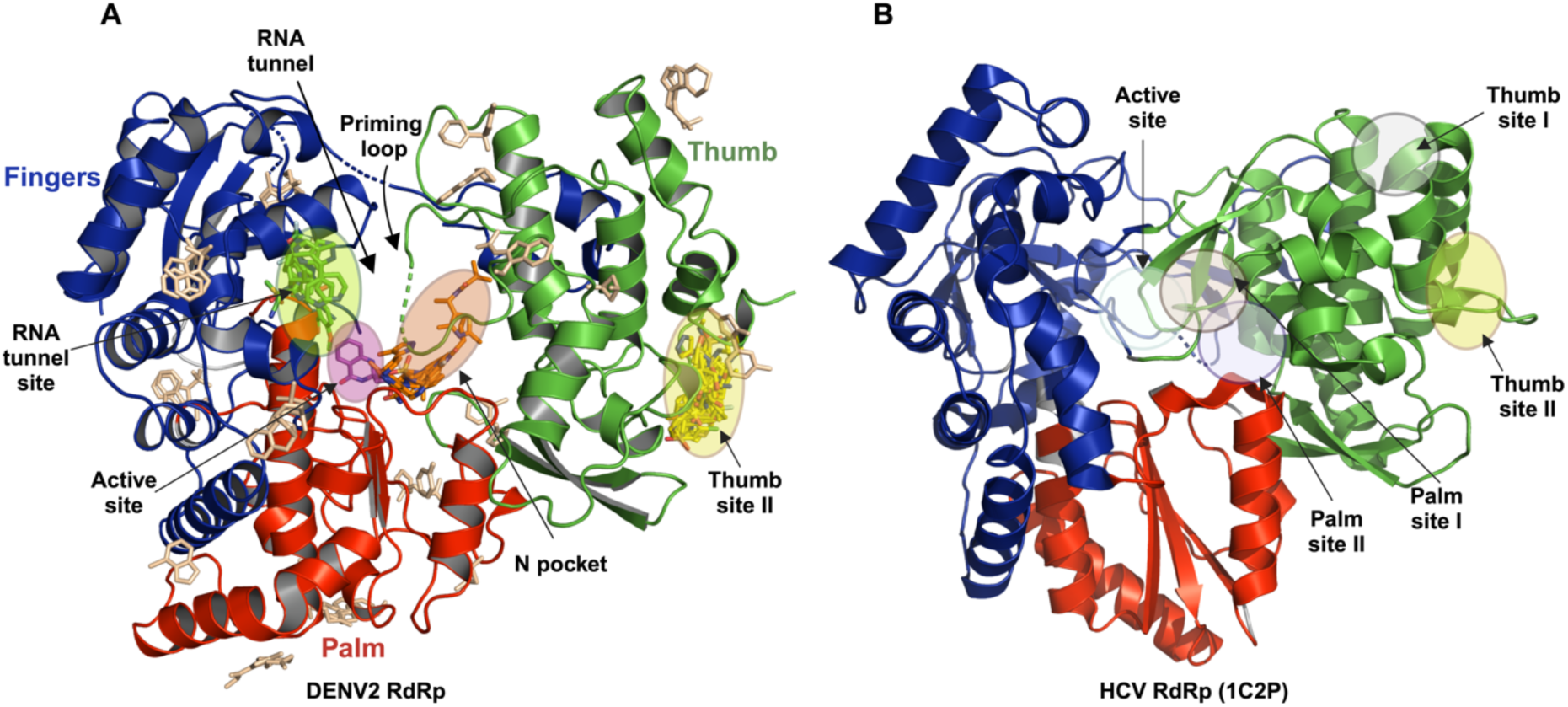
A) Ribbon representation of the overall structure of DENV2 RdRp (residues 273 to 891) with identified fragment hits. The subdomains are color-coded as follows: fingers in blue, palm in red, and thumb in green. B) Comparative side-by-side view of the DENV and HCV RdRp structures, highlighting the active site and other allosteric sites.

First, in this fragment screen we have identified a hit, Z48978335, that binds in the active site of DENV2 RdRp, occupying a spatial region similar to the initiating ATP of the related JEV RdRp^36,37^ and interacting with the GDD conserved catalytic motif (Fig. 2E). This indicates the potential for designing elaborated analogs that can block the initiation step of RNA synthesis. Indeed, flaviviruses have RdRps that initiate RNA synthesis *de novo*, i.e., without the use of a pre-existing primer. Given that the active site is highly conserved across all four dengue serotypes and flaviviruses, such hits have the potential to serve as pan-flaviviral inhibitors.

Regarding the previously targeted N pocket in flaviviruses, it is worthwhile to note that the HCV RdRp palm site I, albeit with a different site architecture, has delivered the FDA-approved drug dasabuvir, and the related compound RG7109 (Fig. 3E and S4B)^51,52^. As mentioned, the retraction of the priming loop from the active site during enzyme elongation may alter the conformation of the N pocket and decrease the affinities of the lead compounds targeting this pocket. Related to this, our fragment hits in the N pocket largely do not engage with the priming loop, in contrast to the previously identified^19^. Therefore, merging moieties from both our fragment hits and from the previous lead series^20^ may be beneficial to preserve the “best of both worlds”: keeping interaction with the priming loop to ‘lock it into place’ to inhibit initiation but also engaging compounds in the area close to the exit dsRNA loop (Fig. 3C and 7A-B). Additionally, the presence of C709 in the primer grip (conserved across all *Flaviviridae* RdRps^19,20^, Fig. S1B) suggests the possibility to design a novel series of covalent molecules utilizing the Z198194394 scaffold as a starting point, which forms an H-bond with this cysteine (Fig. 3Bvi).

Another attractive vector to explore may be extending the N pocket binders to the active site, merging it with hit Z48978335 (Fig. 7E): the engagement with these two regions has the potential to generate tight binding inhibitors. Merging fragments can enhance binding affinity by utilizing multiple interactions between fragments and the protein backbone. Compounds such as Z4628742292, Z55669204, Z198194394, POB0029, and POB0015 (Fig. 3B) can serve as connectors, linking the active site and the N pocket.

The RNA tunnel site has also been previously validated as a viable site across different DENV serotypes^32^. Ligands which bind here appear to interfere with the incoming RNA template and hinder the interaction with the protein backbone during the first steps of RNA polymerization. However, as it is a hydrophobic and relatively enclosed pocket, the fragment and small molecule hits occupy a similar space (Fig. 4). Thus, while there is potential for fragment merging, there is the handicap that they exploit mostly the same limited interactions. As indicated for the N pocket, an intriguing prospect might be the linking of the RNA tunnel site Z1198177230 hit with the aforementioned active site hit (Fig. 7F).

Regarding the thumb site II, we have found that it is the “hottest spot” with 29 fragment hits bound in addition to two previously published compounds, accounting for many merging opportunities. Nevertheless, as a substantial number has crystal packing mediated interactions, priority should be given to the fragments displaying polar interactions with the arginine patch residues involved in RNA recognition, which have been validated as essential for viral replication^45^. In this sense, Fig. 7D lays out the prospect for a manual merge emerging from these “genuine hits”.

The thumb site II is approximately 35 Å from the active site in both HCV and DENV RdRps, and the mechanism by which binding at this region inhibits polymerase activity was elucidated for HCV through a combination of biochemical and biophysical studies with enzyme constructs with deletions or mutations in the beta loop (priming loop notation for HCV) and the C-terminal tail, resulting in a loss of activity in HCV^33,34^. Indeed, these studies observed that inhibitor binding provides stabilization of an inactive RdRp closed conformation of the beta loop, propagated via the thumb site II site. In the case of DENV2 RdRp, we have observed how in several of the thumb site II bound structures the whole priming loop is ordered (contrary to most of the structures of DENV2/3 RdRps, Fig. 6C), suggesting an allosteric relationship between this site and the priming loop.

However, our analysis of the recent Dengue RdRp and NS5 structures reveals another potential mechanism. As mentioned, the thumb site II in DENV RdRp has an arginine patch that was observed to be essential for viral replication^45^. More recently, Osawa and coworkers^13^ have solved cryo-EM structures of NS5 in complex with the flaviviral characteristic 5′-stem-loop structure (SLA). Fig. 6D shows how there is a binding interface between a part of the SLA and thumb site II that overlaps with the fragment hits we have observed in the site. Comparison with the HCV RdRp complex with thumb site II inhibitor filibuvir (as commented, it underwent Phase II clinical trials) shows that in the later the pocket is also highly positively charged, which suggests that in HCV (also with a highly structured 5’ UTR region^53^) such a mechanism could also play a role in the mechanism of inhibition.

In summary, our crystallographic fragment screening campaign has comprehensively charted the binding sites of DENV2 RdRp, that have some commonalities and distinct features to the most targeted RNA virus polymerase, the HCV RdRp NS5B (Fig. 7). Overall, these findings offer a path for developing pan-flaviviral non-nucleoside RdRps inhibitors, a current unmet need in pandemic preparedness.

## Methods

### Expression and purification of the protein

The DENV2 RdRp was codon optimized for *E. coli* expression and synthesized by Genscript in the pET28a vector. The pET28a–DENV2 RdRp construct containing the gene of interest was transformed into *E. coli* BL21 (DE3) star cells and plated on LB agar with 50 μg/mL of kanamycin. A single colony was inoculated in 10 mL of Luria Broth media for primary culture, followed by overnight incubation at 37°C. This primary culture (A_600_= ∼ 1) was used to inoculate one liter LB media (secondary culture). The secondary culture was incubated at 37°C at 180 rpm of shaking until its optical density at 600 nm reached a value of 0.6. The secondary culture was induced with 0.4 mM IPTG (Bio) and incubated at 16°C for 16 h at 180 rpm of shaking.

The cells were harvested at 4000xg and resuspended in a Lysis buffer of 20 mM HEPES pH 7.0, 300 mM NaCl, 10% (v/v) glycerol and EDTA-free protease inhibitor cocktail. IGEPAL CA-630 was then added to a final concentration of 0.1% (v/v), followed by addition of 0.05% (v/v) polyethylenimine (PEI) to precipitate nucleic acid. The lysate was slowly stirred at 4°C for 15 min, sonicated, and then centrifuged for 60 min at 44,000*g* (4°C). The supernatant was purified by nickel-nitrilotriacetic acid (Ni-NTA) affinity chromatography by washing unbound protein with the buffer supplemented with 10 mM imidazole. The RdRp was eluted in a stepwise gradient manner ranging from 40 to 200 mM imidazole. For removing the N-terminal His tag, in house produced HRV14 3C protease was added to the pooled fractions containing the RdRp in 1:50 ratio; the mixture was dialyzed overnight against the buffer-20 mM HEPES pH 7.0, 300 mM NaCl and 1 mM tris(2-carboxyethyl) phosphine (TCEP). The uncut protein pool was separated from the cleaved protein via reverse Ni-NTA. The cleaved RdRp protein was further purified by size exclusion chromatography using 20 mM HEPES pH 7.0, 300 mM NaCl and 5 mM TCEP. SDS-PAGE analysis of the resulting RdRp indicated a purity of ∼95%.

### Crystallization and structure determination

Crystals were grown in a crystallization solution comprising 0.35 M MgCl_2_ along with 0.1 M MES at pH 6.6, and 10% PEG 4,000 maintained at 4°C for one week. An initial model was obtained at a resolution of 1.6 Å (collected at APS beamline 23ID-D), processed within the I222 space group using HKL 2000^54^, and subsequently solved through molecular replacement employing the PDB 5K5M as a reference model with Phaser^55^ and refined with phenix.refine^56^.

For fragment screening, seeds were prepared from crystals that were aspirated and vortexed with 100 μL of the crystallization solution along with glass beads. After optimization, crystals grew reproducibly at 20°C using sitting-drop vapor diffusion in MRC 3 Lens Crystallization plates (SWISSCI) with 300 nl of the aforementioned reservoir solution, with a 1:2 v/v ratio of protein to crystallization solution. This process led to the acquisition of viable crystals in in approximately 80% of the drops within 48 h of the plate setup. To determine solvent tolerability, crystals were incubated with dimethyl sulfoxide (DMSO) ranging 10-20% and incubated for periods of 1-3 h at room temperature.

### Fragment screening

The crystallographic fragment screen was performed using the XChem platform available at Diamond Light Source. Crystal soaking was done by transferring fragments from DSi-Poised Library, Minifrags, Fraglites, Peplites, York3, Halo, SpotXplorer^22,57-59^ to the crystal drop using ECHO Liquid Handler^60^ at proportion of 10% v/v DMSO. Crystals were incubated for 1-3 h at room temperature and harvested using the Crystal Shifter^61^ (Oxford Lab Technologies), mounted on loops and cryo-cooled in liquid nitrogen. All data was collected at I04-1 beamline at Diamond Light Source at 100 K and processed with automated pipelines. All further analysis was performed using the XChemExplorer^62^. Initial density maps were generated using DIMPLE^63^ and with PDB 5K5M (with heteroatoms stripped) as template model. Ligand restraints were generated with ACEDRG^64^ and GRADE^65^. A ground-state model was constructed utilizing PanDDA by amalgamating 100 apo datasets. This model, deposited under the code 7I2X (Table S1), served as the template for molecular replacement for the full analysis. Event maps were calculated with PanDDA^24,66^, and ligands were modeled using Coot^67^. Models were refined with Refmac^68^ using ensemble refinement^68^, and phenix.refine^56^, and models and quality annotations cross-reviewed. Coordinates, structure factors and PanDDA event maps for all data sets are deposited in the Protein Data Bank (PDB, Group deposition IDs G_1002317 and G_1002341). Data collection and refinement statistics, PBD codes and ligand identifier are available in Table S1.

### Reporting summary

Further information on research design is available in the Nature Portfolio Reporting Summary linked to this article.

## Data availability

The coordinates and structure factors have been deposited in the Protein Data Bank. The accession codes are listed in Supplementary Data Table S1. Other data are available from the corresponding authors upon reasonable request. Additionally, the structures have been made available on Fragalysis: https://fragalysis.diamond.ac.uk/viewer/react/preview/target/DENV2_NS5_RdRp/tas/lb32633-1 and https://fragalysis.diamond.ac.uk/viewer/react/preview/target/Flavi_NS5_RdRp/tas/lb32633-1.

## Acknowledgments

The authors would like to acknowledge the Diamond Light Source for access to the fragment screening facility XChem, for usage of DSi-Poised and other libraries and for beamtime on beamline I04–1 under proposal LB32633.

Research reported in this publication was supported by the National Institute of Allergy and Infectious Diseases of the National Institutes of Health under Award Number U19AI171292. The content is solely the responsibility of the authors and does not necessarily represent the official views of the National Institutes of Health.

## Author contributions

EA, FvD, DF and FXR conceived the study and provided supervision. MS carried out protein expression, purification, crystallization, and diffraction data collection. JCA, PGM, BHB and AC contributed to crystal optimization for XChem fragment screening. MS, FXR, and JCA analyzed and processed the data with support from AVC, DF and FvD. AC, and BHB assisted with crystal harvesting and freezing, and data collection. MS, FXR, JCA and DF wrote the manuscript with the assistance of all the authors.

