## Supplementary Data for "Crystallographic fragment screening of the dengue virus polymerase reveals multiple binding sites for the development of non-nucleoside antiflavivirals"

This PDF file includes:

Supplementary Figure 1 to 4

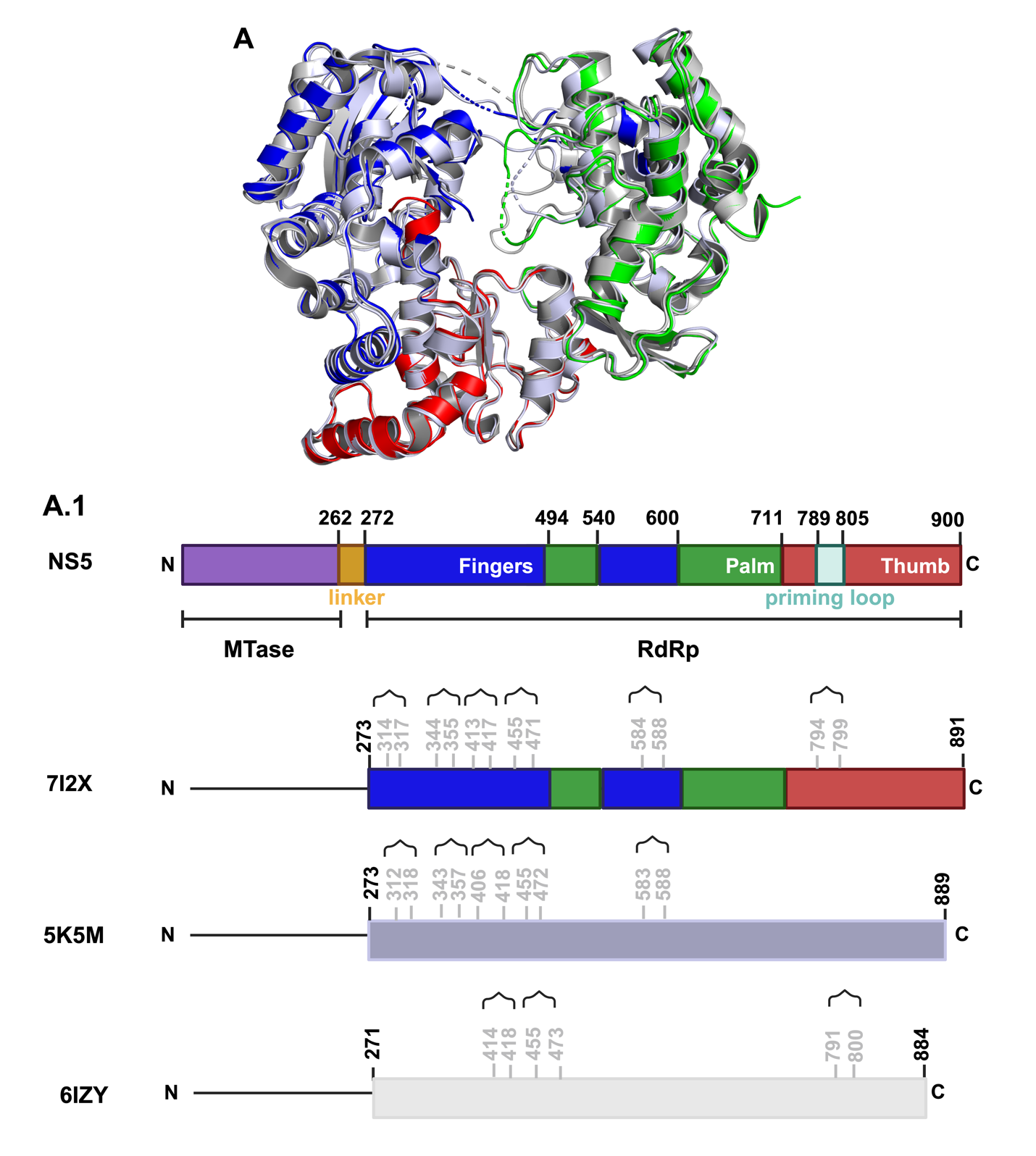

**B**

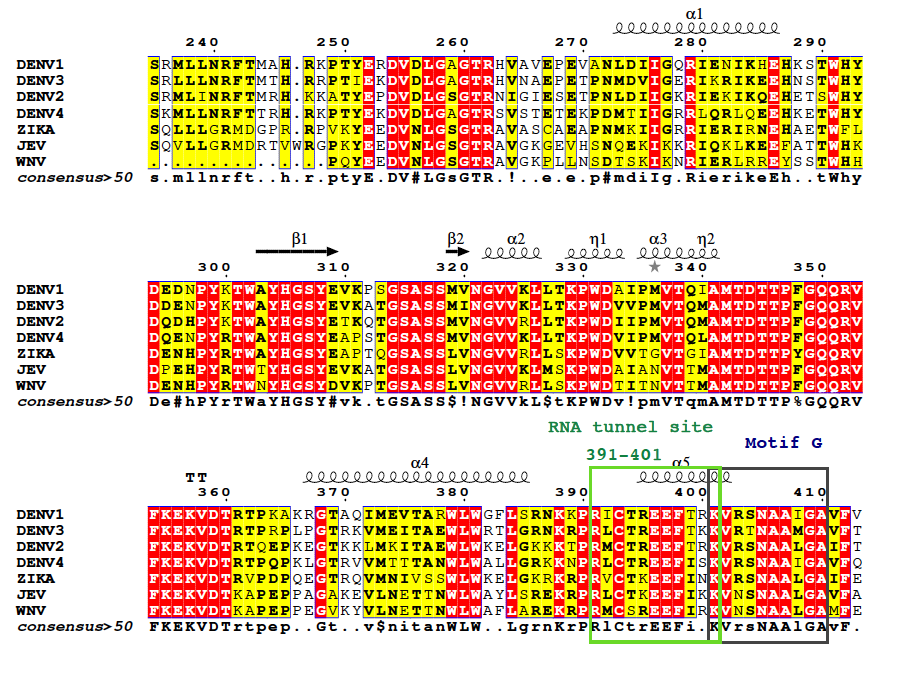

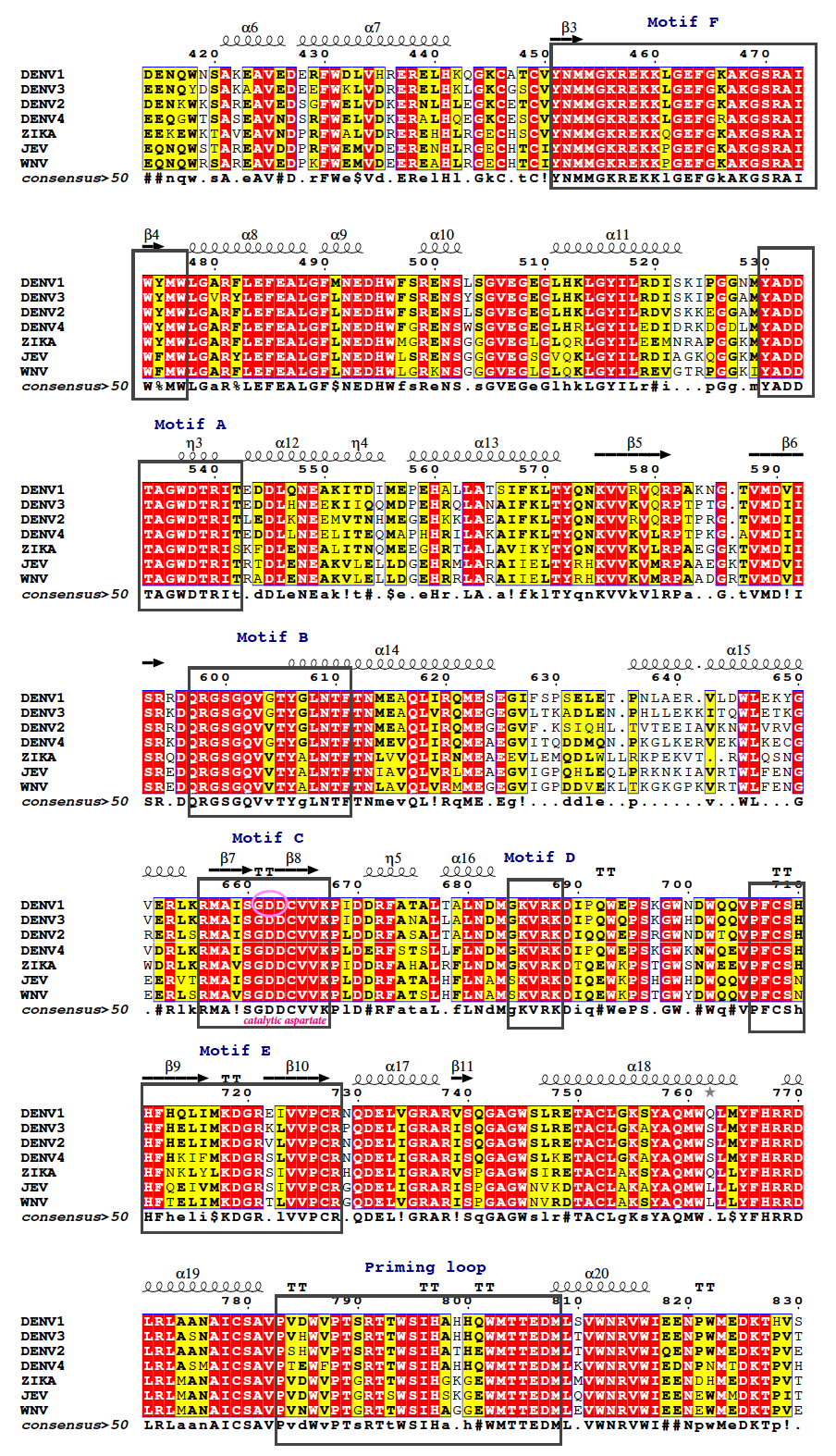

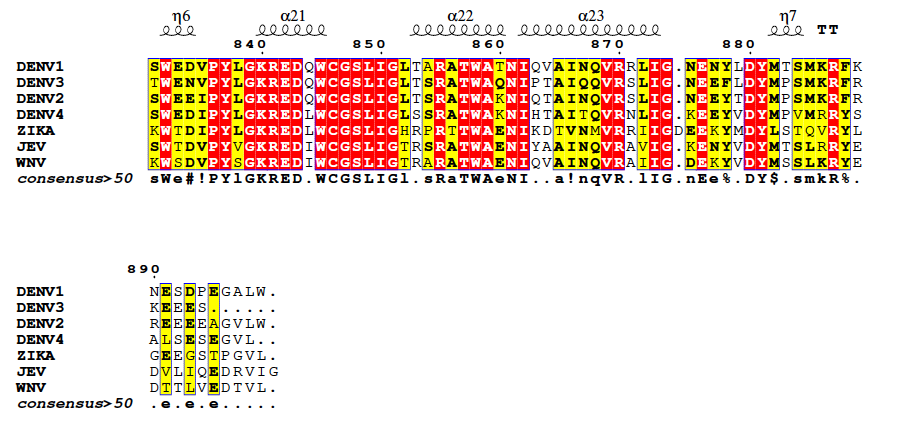

**Figure S1.** Structural comparison of the unbound three-dimensional structure of DENV2 RdRp (PDB: 7I2X) with previously reported structures (PDB: 5K5M and 6IZY). A) The overlay highlights structural similarities and differences, providing insights into conformational variations among these RdRp structures. A.1) Non-observed regions are highlighted in gray. B) Multiple sequence alignment of DENV2 with all its serotypes and close homologs. The conserved structural motifs of RdRps, i.e., A–G, are highlighted in box. The figure was generated using ESPript 3 (<https://espript.ibcp.fr/>).

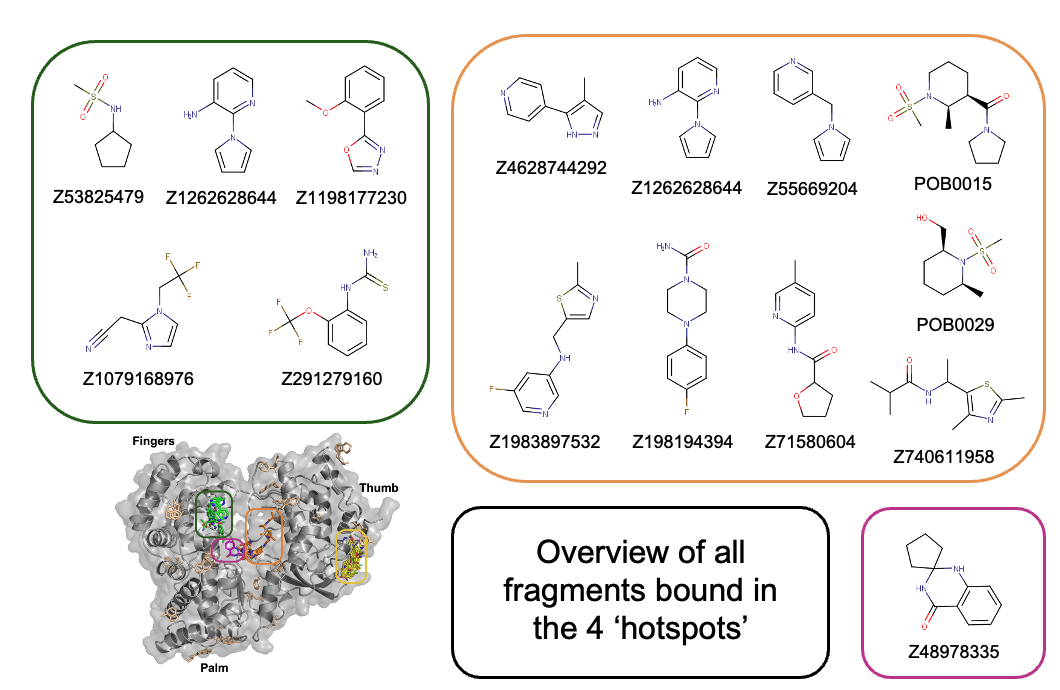

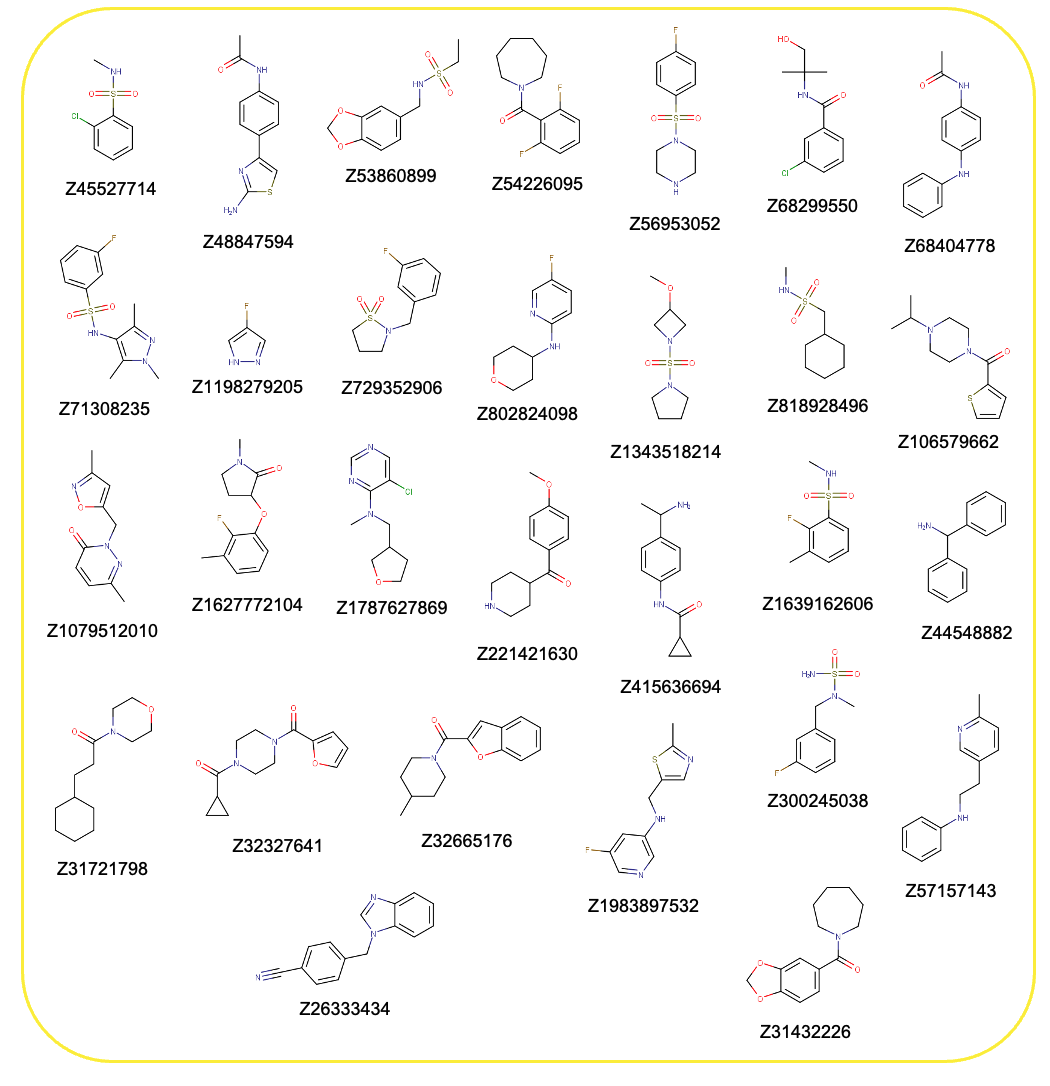

**Figure S2.** Full list of the fragment hits of the four nominated sites. Pink box: Active site; orange box: N pocket or Primer grip site; green box: RNA tunnel site; yellow box: Thumb site II.

**A**

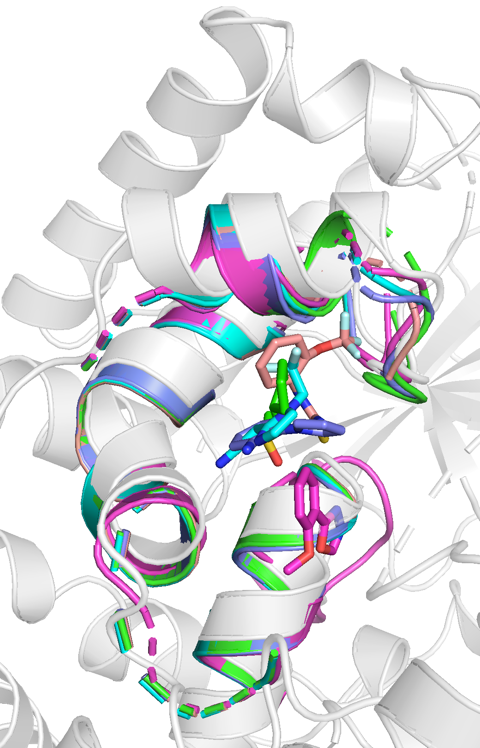

**B**

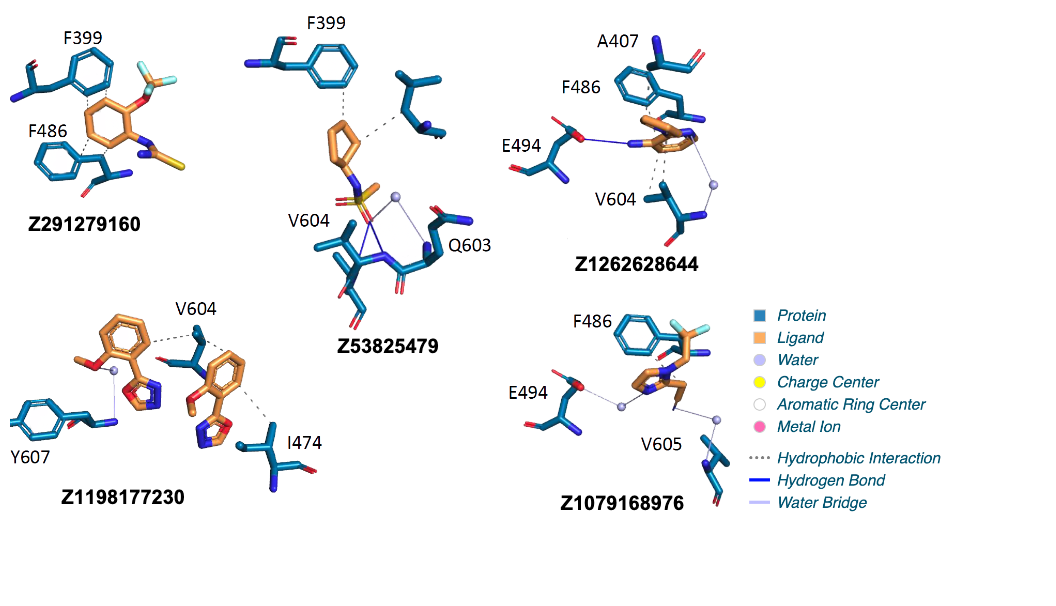

**Figure S3.** DENV2 RdRp RNA tunnel site detailed view. A) Conformational variation of the RNA tunnel site upon fragment binding. Legend: Z53825479 - PDB 7HKD (green), Z1079168976 -PDB 7I2F (cyan), Z1198177230 - PDB 7I2H (pink), Z1262628644 - PDB 7I2I (violet), Z291279160 - PDB 7HKU (light pink). B) Detailed interactions for each fragment hit in the RNA tunnel site, using the PLIP server (<https://plip-tool.biotec.tu-dresden.de/plip-web/plip/index>).

**A**

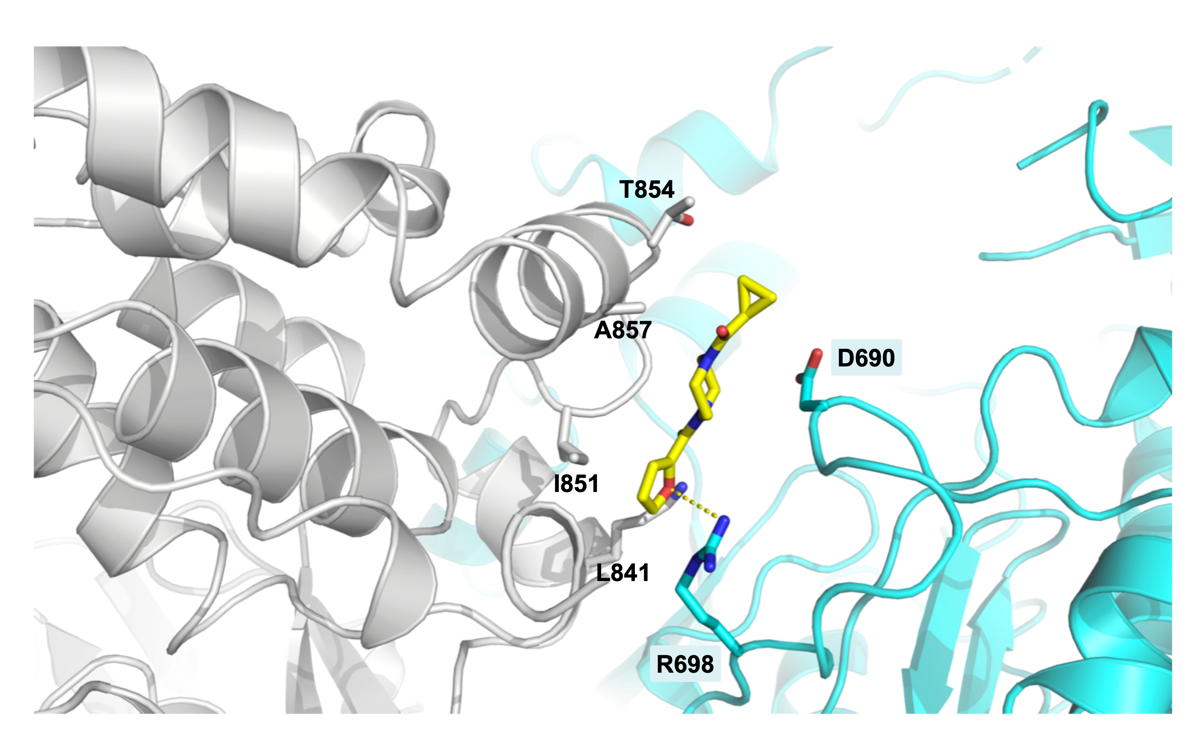

**B**

| **Compound name**  **(and binding site)** | **Structure** | **Pubchem ID** |
| --- | --- | --- |
| **Dasabuvir**  **(Primer grip)** | 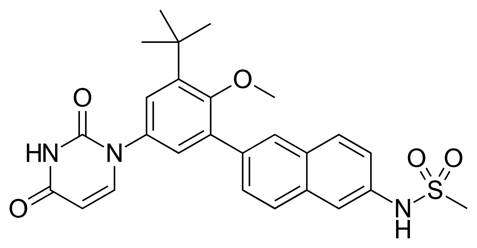 | [56640146](https://pubchem.ncbi.nlm.nih.gov/compound/56640146) |
| **RG7109**  **(Primer grip)** | 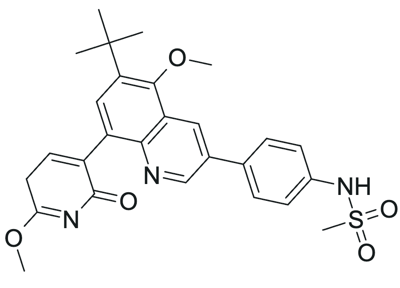 | [137348037](https://pubchem.ncbi.nlm.nih.gov/compound/137348037) |
| **Filibuvir**  **(Thumb site II)** | 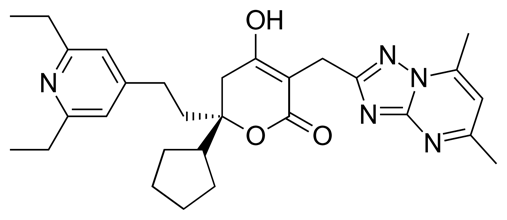 | [54708673](https://pubchem.ncbi.nlm.nih.gov/compound/54708673) |
| **Lomibuvir**  **(Thumb site II)** | 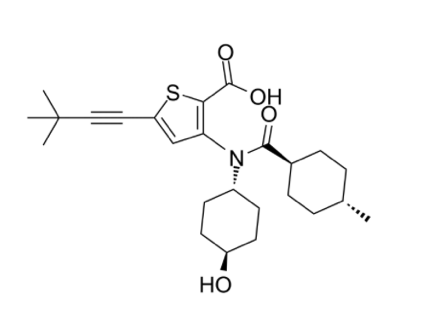 | [24798764](https://pubchem.ncbi.nlm.nih.gov/compound/24798764) |
| **Radalbuvir**  **(Thumb site II)** | 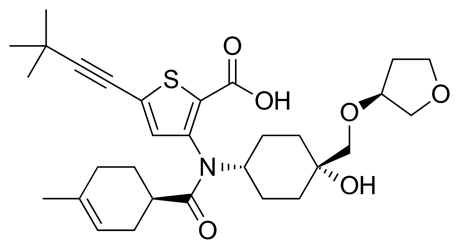 | [53259022](https://pubchem.ncbi.nlm.nih.gov/compound/53259022) |

**Figure S4.** A) Certain binding interactions at the DENV2 RdRp thumb site II were influenced by crystal packing such as with R698 of the neighboring asymmetric unit (cyan). B) Known HCV non-nucleoside inhibitors (NNIs) with their two-dimensional structures and pubchem IDs.
